# SEIR: a novel multi-locus GWAS method that provides higher statistical power for fast identifying variant-phenotype associations

**DOI:** 10.1101/2024.05.23.595530

**Authors:** Guang-liang Zhou, Yun-xia Zhao, Jia-kun Qiao, Fang-jun Xu, Ren-zuo Kuang, Mi-lin Li, Dao-yuan Wang, Ming-yang Hu, Xiao-lei Liu, Xin-yun Li, Shu-hong Zhao, Meng-jin Zhu

## Abstract

Multi-locus genome-wide association study (GWAS) methods have considered the joint effects of multiple variants to more accurately unravel the genetic basis of complex traits. Here, we developed a novel multi-locus GWAS method named Selector-Embedded Iterative Regression (SEIR), which integrates the embedded selector with fast single-marker scanning in an iterative manner. SEIR has excellent adaptability and flexibility under various genetic architectures for qualitative and quantitative traits. Reliability of SEIR was experimentally supported by integrating GWAS with 3D epigenomics in a real trait. Conclusively, SEIR exhibits higher statistical power for fast identifying putative variants compared to other single- and multi-locus methods.

## Introduction

Genome-wide association study (GWAS) has served as a versatile tool for detecting associations between variants and target traits. In GWAS, the individuals are assumed to be unrelated and of the same population background, but this assumption may not hold in practice [1]. Population stratification and cryptic relatedness are common in many current genome-wide association studies and often lead to false positives [2]. To address false positives from population stratification, population structure is typically incorporated as a covariate in general linear modeling (GLM) [3]. However, for individuals with complex and continuous relatedness, that solution is ineffective for eliminating false positives [4]. By contrast, Mixed Linear Models (MLM) can effectively eliminate false positives by simultaneously incorporating these two confounding factors [5]. In MLM, population stratification is fitted as a fixed effect through population structure [6] or principal components [4]. To uncover cryptic relatedness, kinship among individuals is derived from all Single Nucleotide Polymorphisms (SNPs) and can be used to define the variance-covariance structure of individual effects as random effects.

Since MLM was first published, a series of advanced MLM-based methods have been developed [1, 7–10] based on the common feature of single-locus tests, which allows robust analysis of large GWAS datasets. However, for complex traits controlled by multiple, large-effect loci, these approaches may not be appropriate because single-locus test is a one-dimensional genome scan in which one marker is tested at a time and cannot account for the confounding effects of multiple loci. To some extent, this problem is address by multi-locus models, such as Bayesian LASSO [11], stepwise regression [12], penalized or regularized regression [13], and empirical Bayes methods[14]. Although these methods are designed for multi-locus analysis and can be adopted as a model for GWAS, when the number of SNPs (p) exceeds sample size (n), these methods face the problem of p >> n [15] and become computationally unfeasible.

Alternatively, the multi-locus mixed model (MLMM) approach uses simple stepwise mixed-model regression with forward inclusion and backward elimination [16]. Fixed and random model circulating probability unification (FarmCPU) divides MLMM into two parts: a Fixed Effect Model (FEM) and a Random Effect Model (REM), and uses them iteratively [17]. Multi-locus random-SNP-effect mixed linear modeling (MRMLM) applies a single marker random effect mixed linear model (RMLM) as an initial screen, then tests the candidate markers by EM empirical Bayes (EMEB) [18]. Bayesian information and linkage-disequilibrium iteratively nested keyway (BLINK) conducts two FEMs and one filtering process [19], with the first FEM testing all markers; after filtering, remaining markers in the second FEM with the best Bayesian information criteria (BIC) are used as covariates to screen all markers again in the next round of first FEM.

These multi-locus methods share the common feature of dividing GWAS into two analytical steps. The first calculates P-values for all SNPs and selects significant ones based on a threshold; the second selects pseudo-Quantitative Trait Nucleotides (QTNs) from among these significant SNPs. The MLMM method selects pseudo-QTNs using stepwise mixed model regression with forward inclusion and backward elimination. The FarmCPU method adapts a bin approach to select pseudo-QTNs, with REM used to determine the best combination of different bins and number of candidate pseudo-QTNs. The MRMLM method uses a multi-locus model (EMEB) to select pseudo-QTNs. The BLINK method uses the linkage disequilibrium information of significant SNPs to select pseudo-QTNs, with BIC values from FEM determining the number of candidate pseudo-QTNs. In MLMM, FarmCPU, and BLINK, pseudo-QTNs are used as covariates in the statistical model to control false positives, but serve as the final output in MRMLM. Thus, the pseudo-QTN selection strategy can greatly affect the performance of multi-locus GWAS methods.

Here, we propose the Selector-Embedded Iterative Regression (SEIR) algorithm multi-locus GWAS method, which integrates an embedded selector into iterative, single-marker scanning. The selector has excellent adaptability and flexibility, as it can encompass any variable selection method, such as subset selection and penalized regression with variable penalty functions. For single-marker scanning, pseudo-QTNs are used as covariates in a FEM to control for false positives. Single marker scanning substantially reduces the number of markers, allowing the embedded selector to rapidly identify suitable pseudo-QTNs with relatively low computational burden. In each iteration, the selector reselects pseudo-QTNs as covariates in the fixed effect model, and can avoid the problem of some false pseudo-QTNs introduced as covariates in a previous round will be retained in subsequent iterations.

Through GWAS of simulated and real phenotype data from a wide range of species comparing SEIR with other single- and multi-locus GWAS methods, we found that SEIR showed higher statistical power under different False Discovery Rate (FDR) and Type I errors. Furthermore, SEIR could identify as many or more SNPs significantly associated with phenotype through iterative pseudo-QTN selection. Finally, comparison of significant SNPs identified by SEIR with other multi-locus methods in analyses incorporating multi-omics data and 3D genome structure, provides strong proof-of-concept evidence supporting the reliability of associations identified by SEIR. SEIR can thus serve as a powerful and reliable tool for GWAS, with relatively low computational burden, which can be combined with omics phenotype data or 3D genome structure data, to detect genotype-phenotype associations involving multiple genomic loci.

## Results

### Overview of SEIR algorithm

The SEIR method utilizes an iterative approach that integrates single-marker scanning with an embedded selector that includes stepwise regression [12], primal dual active set (PDAS) [20], smoothly clipped absolute deviation (SCAD) [21], minimax concave penalty (MCP) [22], multivariate adaptive regression splines (MARS) [23], least angle regression (LAR) [24], and penalized regression with L1, L0 penalty [25]. In the SEIR framework (Fig. 1a), a FEM is used in the first step to evaluate potential relationships between phenotype and genotype data. After identifying SNPs associated with the trait of interest, a selector is employed to identify pseudo-QTNs for use as covariates in the FEM of the subsequent iteration. This process repeats until no new pseudo-QTNs appear and convergence is reached.

**Fig 1.**
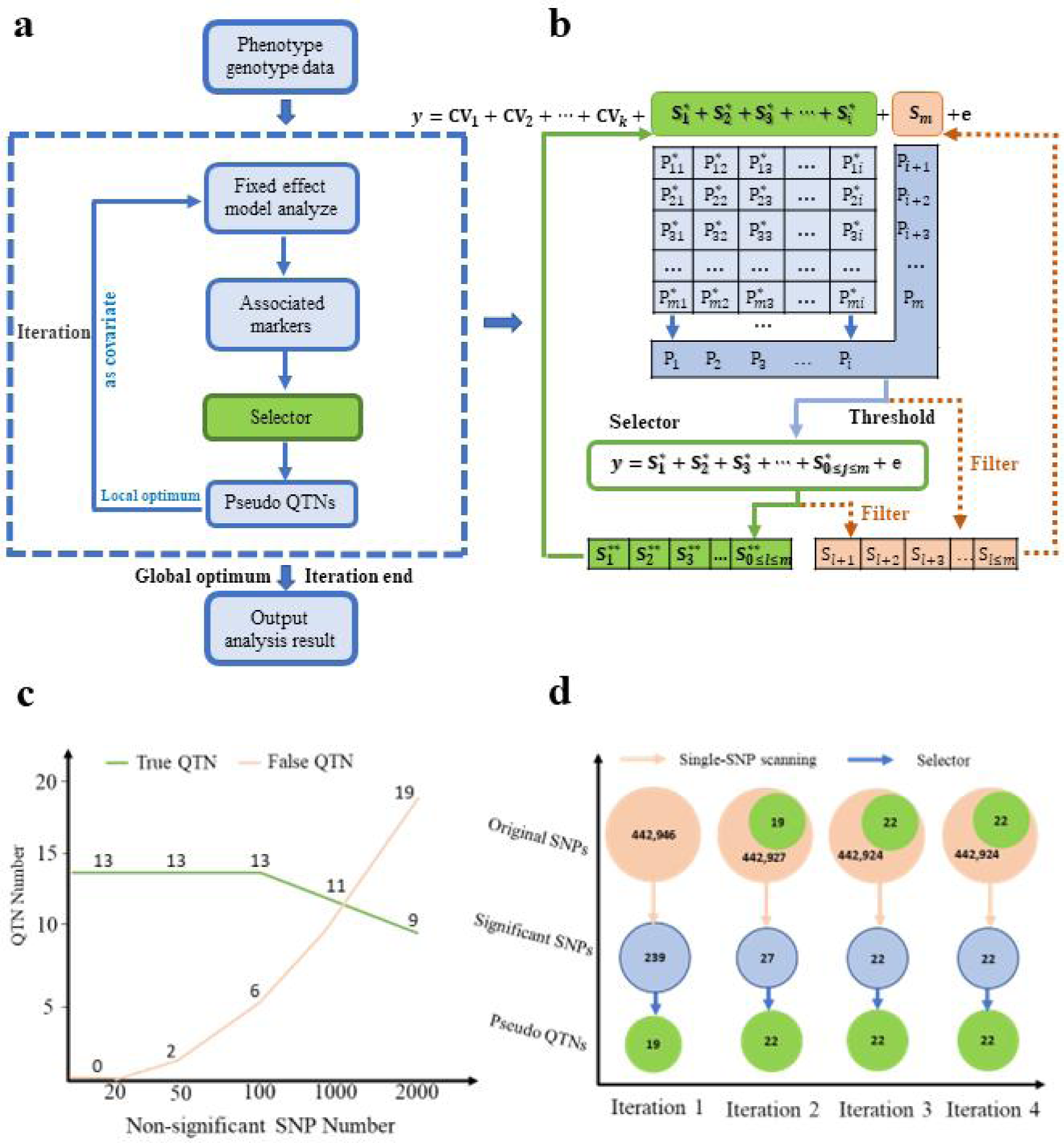
An overview of the SEIR method a Framework of the SEIR method. **b** Detailed process of one iteration in SEIR. The FEM and selector model are used iteratively until no new pseudo-QTNs emerge. CV1 to CVK are other covariates that may influence phenotype. The *i* pseudo-QTNs (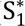 to 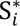) are fitted as covariates to test SNPs one at a time, *e.g*., *m*^th^ SNP (S*m*) in the FEM. As the pseudo-QTNs are fitted as covariates for each SNP, each pseudo-QTN has a test statistic corresponding to every SNP, creating a matrix with elements of 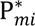, *m* = 1 to m and *i*= 1 to i. The P^∗^value of each pseudo QTN is calculated by the minimum of 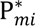 combined with the P-values of remaining SNPs, then filtered. SNPs with P-values larger than a threshold are regarded as pseudo-QTNs (0 ≤ *j* ≤ *m*), while SNPs with P-values below the threshold are regarded as test SNPs (s*_l_*_+1_…s*_l_*_≤*m*_). Pseudo-QTNs are then optimized using one selector (default is stepwise regression). These updated pseudo-QTNs (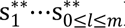) are used as new covariates in the FEM, with the remaining pseudo-QTNs (s*_l_*_+1_…s*_l_*_≤*m*_) also serving as test SNPs. The process repeats continuously until no new pseudo-QTNs are added. **c** The performance of selector in the presence of varying numbers of non-significant SNPs. 20 SNPs were randomly sampled from *Arabidopsis* dataset consists of 1,178 individuals genotyped with 214,545 SNPs, which were used as true QTNs to simulate phenotype. Then, randomly extract another 0, 20, 50, 100, 1000, and 2000 SNPs respectively as non-significant SNPs, and add them to 20 true QTNs to form different genotype data. Using one selector (stepwise regression) to analyze the simulated phenotype with different genotype data. In the absence of non-significant SNPs, this selector can detect 13 true QTNs. Increasing number of non-significant SNPs to 50 or 100, this selector can still detect 13 true QTNs, but also identifies false QTNs. As the number of non-significant SNPs increases, this selector detects fewer true QTNs but a greater number of false QTNs. **d** The detailed process for SEIR to analyze a simulated phenotype. The simulated phenotype with 100 QTNs randomly chosen from a real human dataset. Random error was introduced to fix the heritability at 60%. The dataset includes of 8,807 individuals genotyped with 442,946 SNPs. In each iteration, after single-marker scanning, significant SNPs were filtered by a threshold, and linear dependence is eliminated, allowing the selector to select pseudo QTNs from the remaining significant SNPs with less non-significant SNPs and relatively slight computational burden.

Details of the process of one iteration are shown in Figure 1b. The fixed effect model simultaneously tests *m* SNPs one at a time, and includes pseudo-QTNs as covariates to control for false positives. For each SNP test, each pseudo-QTN has *m* P-values, and the one standard error rule (1se rule) [26] is used to calculate the P-value of each pseudo-QTN. A P-values threshold (Bonferroni correction, α = 0.01) is then used to select significantly associated SNPs, eliminating linear dependence among them. For the remaining candidate SNPs, a selector is used to update the set of pseudo-QTNs that serve as covariates in the FEM of the next iteration. In addition, we tested the ability of selector to detect QTNs and found that the number of true QTNs decreases with increasing number of non-significant SNPs (Fig. 1c). While, initial single-marker scanning using a FEM can screen out most of the non-significant SNPs from the full set of original SNPs, which improves the selector’s performance in pseudo-QTN selection without increasing computational burden (Fig. 1d).

### SEIR outperforms single-locus methods and other multi-locus methods in statistical power

To compare the statistical power of the SEIR method with single-locus and multi-locus methods under different levels of FDR and Type I error, we tested SEIR in a series of simulation scenarios that involved different groups of SNPs representing a variety of genetic architectures. FDR was defined as the proportion of false positives among the total number of positives. Type I error was derived from the empirical null P value distribution of all non-QTN areas. The receiver operating characteristic (ROC) give a description of the relationship between statistical power and FDR or Type I error. The method with larger area under the curve (AUC) have higher statistical power than the method with a smaller AUC. Phenotypes were simulated by adding the additive genetic effect values and residual effect values, as described in Materials and Methods section.

The single-locus GWAS methods include classical GLM and MLM, as well as some modified MLM methods. Therefore, we first compared SEIR with GLM and MLM using simulated traits with a heritability of 0.6 controlled by 20 or 100 QTNs. SEIR showed higher statistical power than GLM and MLM under different levels of FDR and Type I error (Fig. 3). The comparisons were then extended to simulated traits with different levels of heritability (0.2, 0.8), which showed that SEIR consistently outperformed GLM and MLM (Additional file 1: Fig. S1 and Additional file 2: Fig. S2). Comparison of SEIR with two modified MLM methods, GRAMMAR-Gamma and BOLT-LMM, in the same simulated scenarios and datasets revealed that SEIR could also provide higher statistical power versus FDR and Type I error (Additional file 3: Fig. S3 and Additional file 4: Fig. S4). These comparisons of SEIR with GLM and MLM aligned well previous studies that showed multi-locus approaches could provide more robust QTN identification than single-locus approaches [16–19].

**Figure 2.**
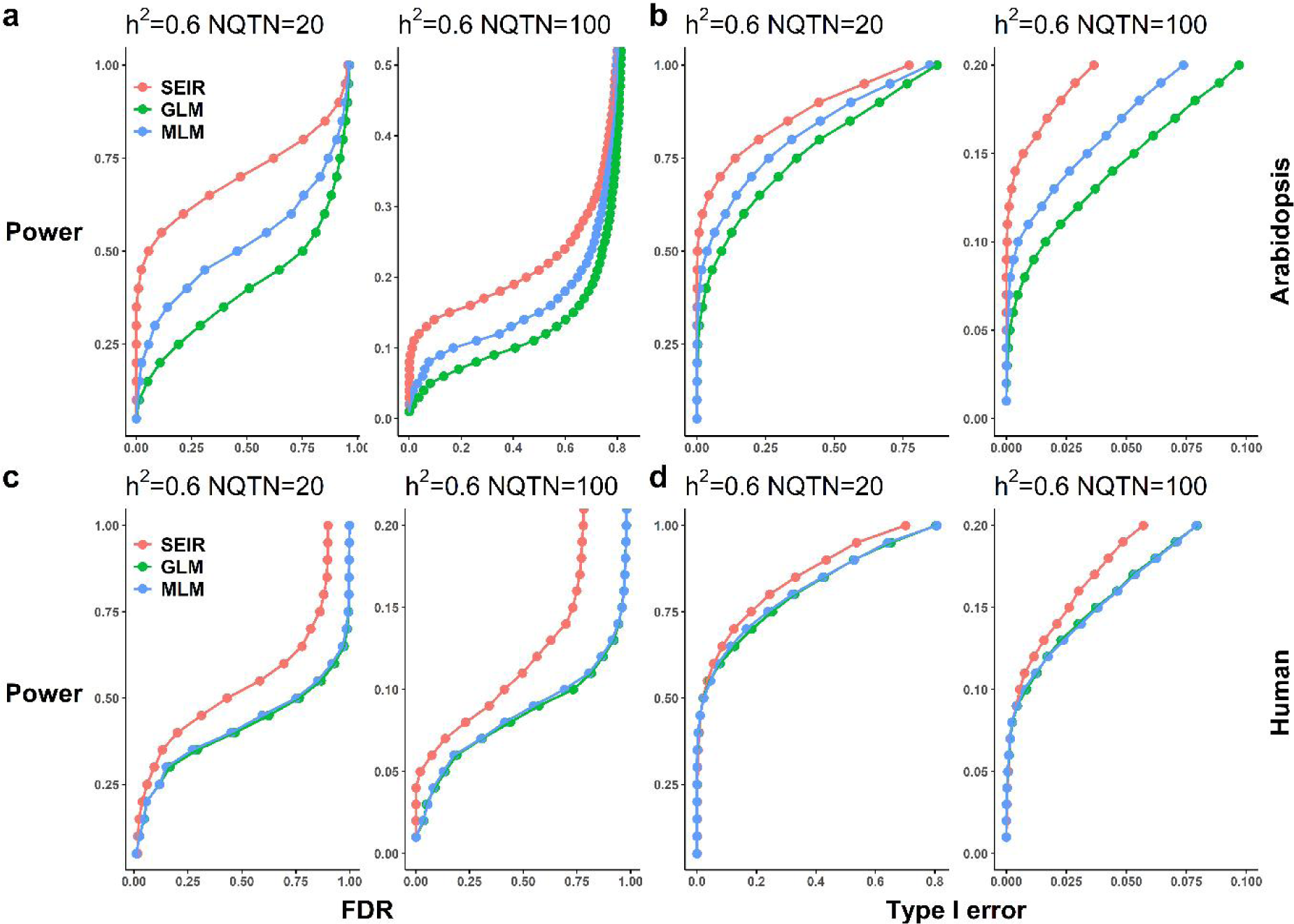
Statistical power of SEIR, GLM, and MLM in different simulation scenarios. Comparison between SEIR and GLM, MLM in different simulation scenarios. The top panel and bottom panel display the simulation results, based on Arabidopsis and human data, respectively. The Arabidopsis dataset consists of 1,178 individuals genotyped with 250,000 SNPs and the human dataset consists of 8,807 individuals genotyped with 442,946 SNPs. Additive genetic effects were simulated with 20 QTNs with normal distribution and 100 QTNs with geometric distribution. The QTNs were randomly sampled from all the SNPs in each dataset. Residuals with normal distribution were added to the genetic effect to form phenotypes with heritability of 0.6. The QTNs were randomly sampled from all the SNPs in each dataset. Power was examined under different levels of FDR and Type I error. A positive SNP is considered a true positive if a QTN is within a distance of 10,000 base pairs on either side; otherwise, it is considered as a false positive. The two types of receiver operating characteristic (ROC) curves are displayed separately for FDR (a, c) and Type I error (b, d)

**Figure 3.**
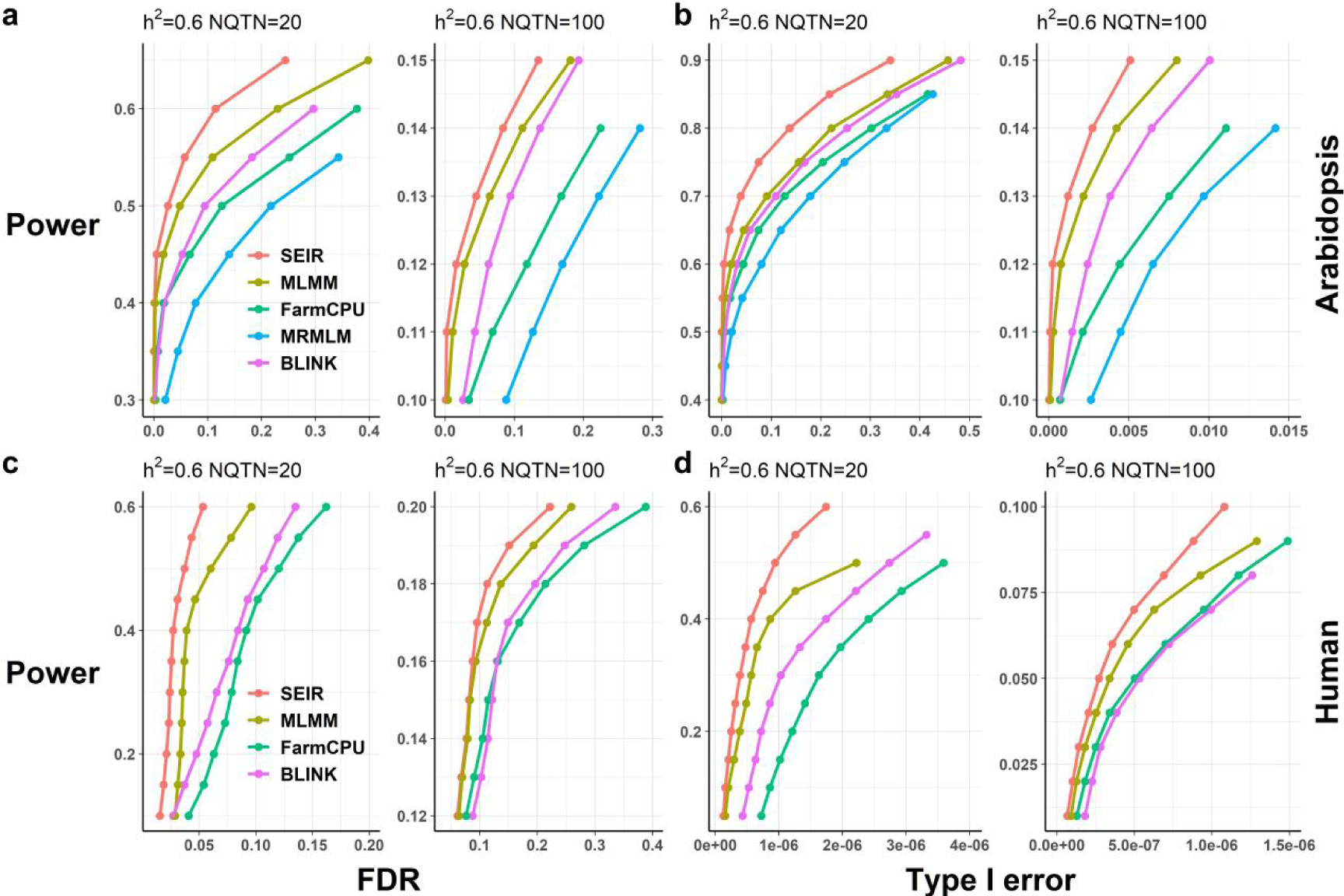
Statistical power of SEIR, MLMM, FarmCPU, MRMLM and BLINK in different simulation scenarios. Compared SEIR with MLMM, FarmCPU, MRMLM, and BLINK in different simulation scenarios. The top panel and bottom panel display the simulation results, based on Arabidopsis and human data, respectively. The dataset from Arabidopsis population consists of 1,178 individuals genotyped with 250,000 SNPs. The dataset from human population consists of 8,807 individuals genotyped with 442,946 SNPs. In particular, there is no MRMLM in the comparison of human data simulation, due to the large size of human data, MRMLM cannot calculate. Additive genetic effects were simulated with 20 QTNs effect with normally distributed and 100 QTNs effect with geometric distribution. Residuals with normal distribution were added to the genetic effect to form phenotypes with heritability of 0.6. The QTNs were randomly sampled from all the SNPs in each dataset. Power was examined under different levels of FDR and Type I error. A positive SNP is considered a true positive if a QTN is within a distance of 10,000 base pairs on either side, otherwise is considered a false positive. The two types of receiver operating characteristic (ROC) curves are displayed separately for FDR (**a, c**) and Type I error (**b, d**)

Multi-locus GWAS methods such as MLMM, FarmCPU, BLINK, and SEIR adopt different strategies for identifying pseudo-QTNs that could potentially result in selecting different pseudo-QTNs as covariates to control for false positives in the statistical model, and consequently, variation between methods. As a result, the statistical power of these methods may vary. We therefore compared SEIR with MLMM, FarmCPU, MRMLM, and BLINK using the same scenarios and publicly available data as in the simulations above. These analyses showed that SEIR consistently provided higher statistical power than the other four methods under different levels of FDR and Type I error, at simulated traits with heritability of 0.6 controlled by 20 or 100 QTNs (Fig. 3). Similar results were obtained for simulated traits with heritability of 0.2 or 0.8 controlled by 20 or 100 QTNs (Additional file 5: Fig. S5 and Additional file 6: Fig. S6). In addition, we assessed the performance of these multi-locus methods in complex traits with heritability of 0.6 controlled by 500 or 1000 QTNs, which further supported that SEIR could provide higher statistical power than the other four methods (Additional file 5: Fig. S5 and Additional file 6: Fig. S6).

### The pseudo-QTN selection by SEIR identifies more true QTNs than other multi-locus methods

Differences in pseudo-QTN selection among multi-locus methods can lead to differences in the above statistical power comparison for each approach. While SEIR, MLMM, FarmCPU, and BLINK iteratively select pseudo-QTNs from the associated SNPs, MRMLM employs EMEB for only a single round of pseudo-QTN identification. Thus, for compare the differences between different multi-locus methods in pseudo-QTN selection, the associated SNPs identify by SEIR with those detected by other methods using the same datasets as for the above phenotype simulations were further analysis.

For this analysis, we used a complex phenotype scenario with heritability of 0.6 controlled by 500 QTNs in the Arabidopsis genotype data containing 250,000 SNPs from 1,178 individuals. Since QTN locations were known in this simulation, we compared the distance between these QTNs and the associated SNPs identified by each multi-locus method. Associated SNPs for each method were defined as those with P-values below the threshold (Bonferroni correction, α = 0.01). Distances between an associated SNP and true QTNs were divided into four categories in which 0 kb indicated a match, 0<10 or 10<50 kb on either side of the QTN suggested potential association, and distances >50 kb were considered spurious associations.

The results revealed that a significantly higher proportion of associated SNPs identified by SEIR were true QTNs compared to the success rate of other multi-locus methods (Fig. 4a). More specifically, 87% of the associated SNPs were true QTNs, whereas 80% of associated SNPs were true QTNs in BLINK analysis, 78% in FarmCPU, 82% in MLMM, and 19% in MRMLM analyses. By contrast, SEIR found fewer potentially associated SNPs at 0-10 kb distance from the QTN (8%), compared to MRMLM (52%), BLINK (11%), FarmCPU (9%), and MLMM (13%). Among potential associated SNPs at 10-50 kb distance, SEIR had 2.6%, which was lower than MRMLM (22.5%), BLINK (3.4%), and FarmCPU (6.5%), but comparable to MLMM (2.7%). Comparison of spurious SNPs >50 kb distance from a true QTN, MLMM had the lowest proportion (1.2%), while SEIR had 1.8%, BLINK had 4.2%, FarmCPU had 5.5%, and MRMLM had a proportion of 6%.

**Fig 4.**
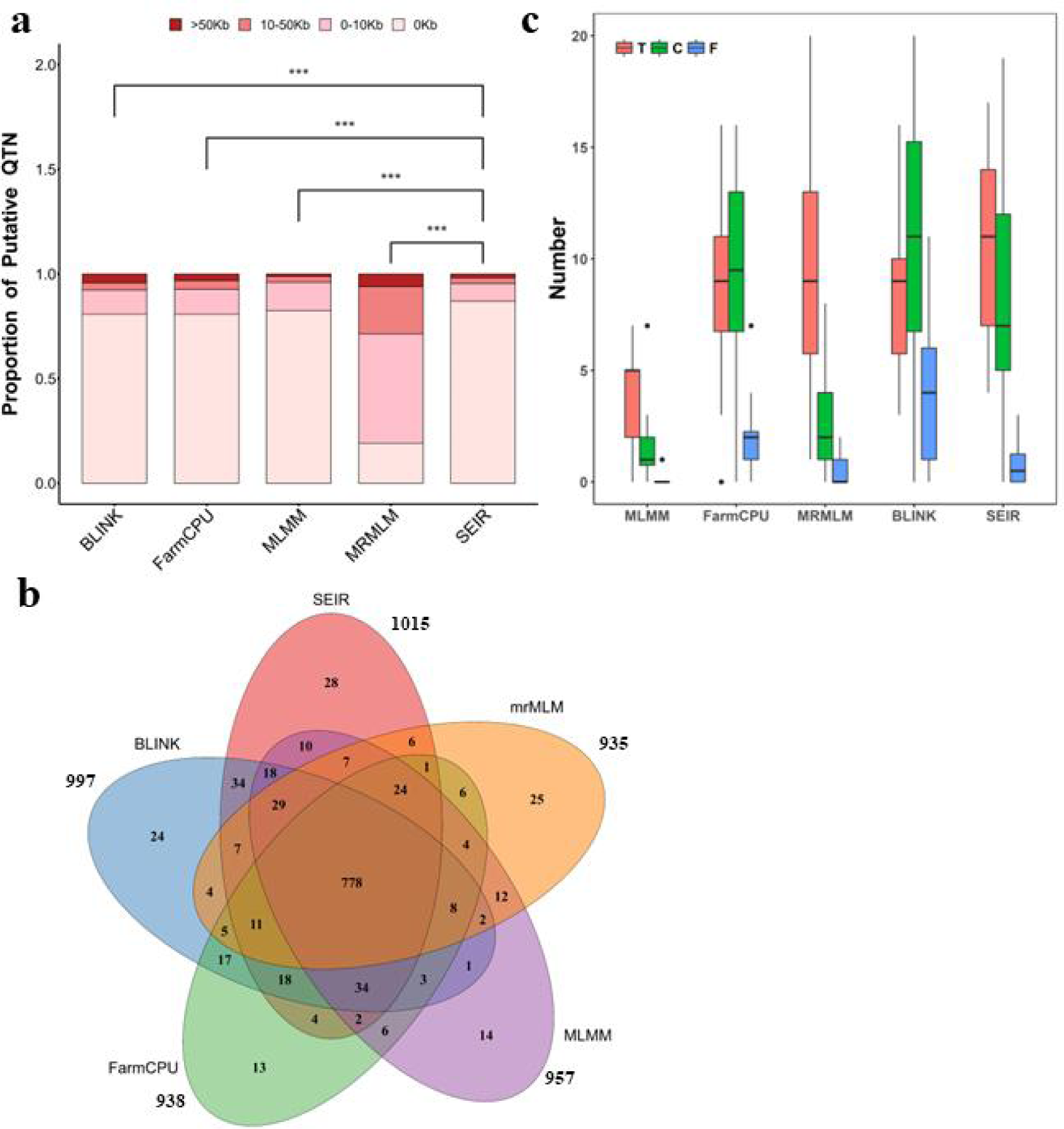
Summary of the associated SNPs detected by SEIR, MLMM, FarmCPU, MRMLM, and BLINK in the simulated and real phenotypes. The simulated complex phenotype includes 500 QTNs randomly chosen from Arabidopsis dataset which was consists of 1,178 individuals genotyped with 250,000 SNPs. The effect of QTNs follows geometric distribution. Residuals with normal distribution were added to the genetic effect to form phenotypes with heritability of 0.6. The simulations were repeated 100 times. Real phenotypes were 20 phenotypes of yeast. **a** The distribution proportion of distance between associated SNPs and QTN detected by SEIR, MLMM, FarmCPU, MRMLM, and BLINK. The distance between associated SNPs and QTN is divided into four categories: 0 kilobase (kb), 0-10 kb, 10-50 kb, and out of 50 kb. Associated SNPs detected by SEIR and other multi-locus methods were significantly different among the four categories. **b** The Venn diagram between the number of QTN identified by SEIR, MLMM, FarmCPU, MRMLM, and BLINK. SEIR identified 1,015 QTNs, higher than MLMM (957), FarmCPU (938), MRMLM (935), and BLINK (997). 778 QTNs were identified by all the multi-locus methods. SEIR identified 28 unique QTNs, higher than MLMM (14), FarmCPU (13), MRMLM (25), and BLINK (24). **c** Statistics results of the match information for SEIR, MLMM, FarmCPU, MRMLM, and BLINK in yeast data. The associated SNPs identified by each multi-locus method were compared with the detailed QTL information to count matching cases. T, C, and F represent the following cases: associated SNPs in GWAS results were consistent with QTL information, within the confidence interval of QTL information, and not in the QTL information. The vertical axis indicates the number of associated SNPs in three cases (T, C, and F) for each method. For each box, the inside line represents the average number of associated SNPs in three cases (T, C, and F), the top and bottom whiskers are the maximum, minimum numbers of associated SNPs in three cases (T, C, and F), respectively. The points outside the box indicate abnormal behavior of a method in a specific trait.

Further analysis revealed that SEIR identified more QTNs at 0 kb from associated SNPs than the other multi-locus methods (Fig. 4b). Among the associated SNPs identified, 778 were identical, and SEIR identified 28 unique associated SNPs that matched QTNs, which were higher than MLMM (14), FarmCPU (13), MRMLM (25), and BLINK (24). The enhanced ability to capture associated SNPs by SEIR primarily reflects the embedded selector approach to pseudo-QTN selection compared to the strategies used by MLMM, FarmCPU, MRMLM, and BLINK.

Additionally, we further verified these findings using genomic data from yeast segregants strain with 20 growth traits that was extensively analyzed in previous work [27], and thus included information on quantitative trait loci (QTL), such as physical locations of QTNs and confidence interval sizes (Additional file 14: Table S1). We re-conducted GWAS for 20 growth traits using SEIR, MLMM, FarmCPU, MRMLM, and BLINK, and compared the associated SNPs identified by each method with the detailed QTL information to determine the accuracy and sensitivity of each method (Fig. 4c). In this analysis, associated SNPs were categorized as either matching a QTN (T), within the confidence interval of the QTN (C), or outside the confidence interval of the QTN (F).

Comparison of associated SNPs in each category indicated that SEIR could identify more associated SNPs that matched QTNs than the other four multi-locus methods. By contrast, BLINK identified the highest number of significant SNPs within the confidence interval of the QTN. While MLMM had the lowest detection rate in all categories. These results also indicated that the differences in pseudo-QTN selection among multi-locus methods can lead to differences in the number of associated SNPs identified by each approach. SEIR could identify more true QTNs with fewer spurious association SNPs than other multi-locus methods.

### SEIR identifies more significant SNPs and fewer spurious associations with real phenotype data compared to other methods

In light of the above results obtained in simulations, we next conducted GWAS with SEIR, MLMM, FarmCPU, MRMLM, and BLINK in genomic data from humans, pigs, chickens, mice, and horses, as well as the model plant, *A. thaliana*, that corresponded to real phenotypes (Additional file 7: Fig. S7-Additional file 12: Fig. S12). We found that some of the associated SNPs identified by these methods were previously reported as associated with a specific phenotype. In lung cancer in Asian never-smoking women, all methods could identify known associated SNPs on chromosomes 5 and 10 [28]. In chicken, neither MLMM nor MRMLM could detect associated SNPs, whereas SEIR, BLINK, and FarmCPU all identified a known putative SNP on chromosome 1 [29]. In horses, all methods could identify a known coat color locus [30]. In Arabidopsis, SEIR could identify a SNP on chromosome 5 significantly associated with flowering time phenotype and matching the known *FLOWERING LOCUS C* (*FLC*) gene [31].

Furthermore, these multi-locus methods could also identify previously unreported significant SNPs. For example, SEIR and BLINK identified the same SNPs on chromosomes 18 and 20 associated with the lung cancer in Asian never-smoking women. Each method also identified unique associated SNPs not detected by other methods, such as the flowering time-associated SNP in Arabidopsis or a SNP associated with backfat thickness in pig. We therefore sought to determine whether associated SNPs among these new findings were true QTNs or spurious associations. As previous studies have shown that permutation tests can effectively screen out spurious associations [32], we used permutation tests to screen the associated SNPs identified by SEIR, MLMM, FarmCPU, MRMLM, and BLINK. This analysis showed that some SNPs were indeed non-significant associations, although notably, SEIR had fewer such spurious associations than that in the other four methods (Additional file 15: Table S2). These cumulative results illustrated that SEIR could identify SNPs at the same, known, phenotype-associated loci as other multi-locus methods while also detecting previously overlooked significant SNPs, and with fewer spurious SNPs.

### Multi-omics data verification of SNP-phenotype associations in a pig data

Based on permutation tests showing that some SNPs identified in the above comparison of GWAS methods using real phenotypes were indeed significantly associated with phenotype, we next examined these significant SNPs with additional omics data, such as epigenetic data. In particular, we focused on GWAS results of SEIR, MLMM, FarmCPU, MRMLM and BLINK screening for SNPs associated with backfat thickness in a study population of 4,555 individual pigs genotyped with 47,258 SNPs.

Several significant SNPs were identified by multiple methods, such as those on chromosomes 1 (MLMM, BLINK, FarmCPU SEIR), 3 (FarmCPU, BLINK, SEIR), and 13 (FarmCPU, MRMLM, BLINK, SEIR) (Fig. 5a). Intriguingly, four methods identified DRGA0007123 as a significant SNP on chromosome 7. Similarly, while DRGA007128 was only detected by BLINK. In addition, linkage disequilibrium (LD) analysis indicated that five SNPs in this region were tightly linked, forming an LD block (chr7: 9912549-10010359) that harbored both DRGA0007123 and DRGA007128. In GWAS, when limited by low marker density, candidate genes may be located based on linkage between SNPs and causal loci. Although previous studies have shown that GWAS combined with three-dimensional genomic data can be an effective strategy for locating candidate genes [33], the relationship between the LD of SNPs and 3D genome structure has not been investigated.

**Fig 5.**
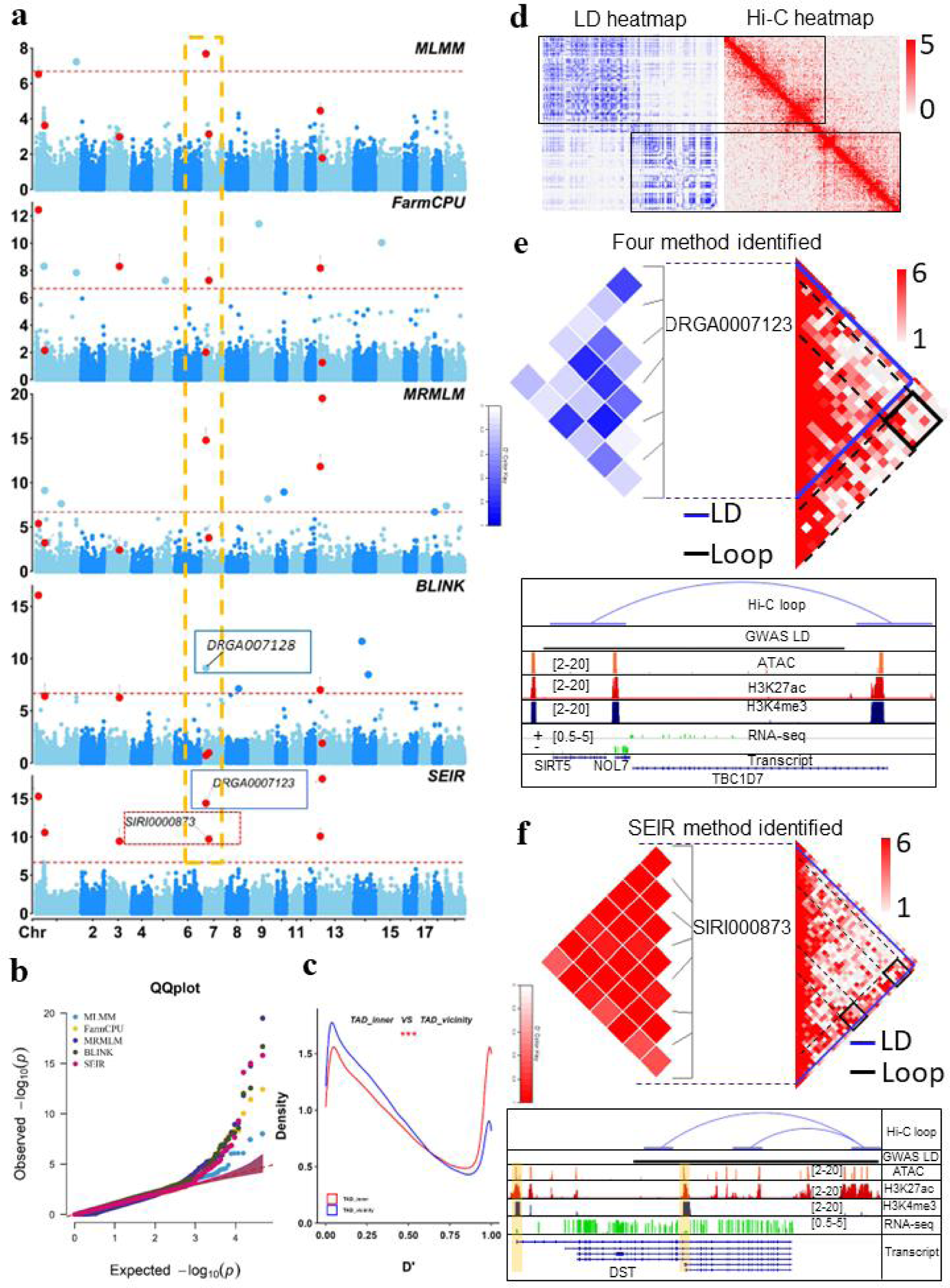
Integrating 3D genome data with GWAS to verify SNP association with backfat thickness. **a** Manhattan plots comparing results of GWAS conducted with MLMM, FarmCPU, MRMLM, BLINK, or SEIR. Red dots, significant SNPs detected by SEIR and other methods; yellow box, significant SNP on chromosome 7 detected by all methods; blue box, DRGA0007123 detected by MLMM, Farm CPU, MRMLM and SEIR; red box, SIRI0000873 detected only by SEIR. **b** QQ plot for MLMM, Farm CPU, MRMLM, BLINK, and SEIR. **c** D’ values of SNPs within the same or neighboring TADs. Red line, density distribution of D’ values for SNPs within TADs (TAD_inner); blue line, density distribution of D’ values for SNPs in vicinity TADs (TAD_vicinity). **d** LD heatmap based on whole-genome sequencing data of Duroc, Landrace, Yorkshire from ISWINE database, and Hi-C contact map of LW pig backfat tissue at 5 kb resolution for the pig genome. **e** Top: LD block (chr7: 9912549-10010359) harboring DRGA0007123 identified by MLMM, Farm CPU, MRMLM and SEIR and DRGA007128 identified by BLINK. Bottom: Tracks for epigenetic modification signals and 3D genome structures around the LD block harboring DRGA0007123 and DRGA007128. **f** Top: LD block (chr7: 29166478-29268803) around SIRI0000873 (chr7: 29228734). Bottom: Tracks for epigenetic modification signals and 3D genome structures around LD block harboring SIRI0000873 identified by SEIR.

Next, the statistics results showed the D’ values of SNPs within the same topologically associating domain (TAD) were significantly higher than those of SNPs in neighboring TADs (Fig. 5c). Furthermore, ∼80% of LD blocks identified in this pig population were located within TAD regions (Additional file 13: Fig. S13). Moreover, a LD heat map generated from whole-genome resequencing data was largely concordant with a Hi-C contact map at 5 kb resolution (Fig. 5d), which supporting LD block regions are strongly associated with 3D genome structure.

Notably, the LD block (chr7: 9912549-10010359) that harbored significant SNP, DRGA0007123 and DRGA007128, as well as, which almost fully covered a Hi-C loop located in a backfat tissue TAD (chr7:9240002-10160000; Fig. 5e). By integrating ChIP-seq and ATAC-seq data from back fat tissue, we revealed this Hi-C loop represented interaction between the *NOL7* gene promoter and the promoter region of the *TBC1D7* gene. RNA-seq data further confirmed that both *NOL7* and *TBC1D7* were expressed in backfat tissue. Closer examination of previous research on *NOL7* and *TBC1D7* indicated that the former was involved in regulating cell growth, while the latter can reportedly regulate *mTORC1* to modulate lipid metabolism [34], aligning well with a potential role in porcine backfat phenotype. These results showed that integrating GWAS with multi-omics data can provide an effective strategy for linking candidate loci with phenotype.

After investigating this common SNP in detail, we next investigated SNP SIRI0000873 chr7: 29228734, which was uniquely identified by SEIR as associated with backfat, and formed an LD block (chr7: 29166478-29268803) with surrounding SNPs. By integrating GWAS results with 3D genome structure data, as in the strategy above, we identified the chr7: 29166478-29268803 LD block containing two Hi-C loop structures in a fat tissue TAD (chr7:28600002-29680000). Remarkably, only one gene, *DST,* was found in this region, the promoter of which was located in an anchor region of the chr7: 29175000-29375000 loop. RNA-seq data indicated that this gene was also highly expressed in adipose tissue (Fig. 5f). These results thus confirmed that the significant SNP identified by SEIR was likely associated with backfat phenotype in pigs, and further illustrated the informative value of combining GWAS with other multi-omics data to verify SNP-phenotype associations.

## Discussion

GWAS methods based on single-locus tests, along with polygenic background and population structure controls, have been used to identify associations between genomic variants and traits of interest in multiple species. However, as noted by Segura et al. [16], single-locus tests risk using the wrong model unless the trait is indeed controlled by a single locus, since these tests fail to account for possible joint effects of multiple genes on phenotype due to intrinsic limitations of the model. Although subset selection and penalized regression with variable penalty functions provide an alternative for multi-locus GWAS, these approaches are challenged by the *p >> n* problem and are computationally unfeasible when the number of markers is much larger than the sample size. Typically, a trait is only associated with a small subset of SNPs, and therefore reducing the number of candidate SNPs before employing a shrinkage method in a dimension-reduced model may alleviate computational burden.

Existing multi-locus GWAS methods typically divide GWAS into two analytical steps. The first step involves screening associated SNPs from a very large number of original SNPs through a variety of statistical models. However, the introduction of spurious associations is a common artifact of screening large SNP datasets. Thus, the second step in many multi-locus methods involves selection of pseudo-QTNs from among the total associated SNPs. However, at least two issues with this second step have emerged: calculating P-values for pseudo-QTNs and selecting appropriate pseudo-QTNs.

To calculate the P-value of pseudo-QTNs, MLMM calculates P-value using all pseudo QTNs as covariates in the model and but excludes testing marker [16]. By contrast, MRMLM calculates the P-value of pseudo-QTNs in EMEB modelling, while SEIR, uses the most significant P-value out of each pseudo QTN in conjunction with the tests on all markers, similar to strategies employed by FarmCPU [17] and BLINK [19]. FarmCPU and BLINK test pseudo-QTNs individually with all markers and select those with the lowest P-values [26], while SEIR incorporates the “1se rule” for P-value-based pseudo-QTN selection. Results in this current study support that use of the 1se rule can improve statistical power to some extent.

To determine the optimal set of pseudo-QTNs, MLMM employs stepwise mixed-model regression with forward inclusion and backward elimination. However, this strategy imposes a heavy computational burden for large sample sizes. In FarmCPU, pseudo-QTNs are selected through binning, but only one pseudo-QTN can be selected as a covariate regardless of whether the bin includes multiple pseudo-QTNs, consequently limiting the statistical power of this method. In MRMLM, the RMLM selects pseudo-QTNs using less stringent criteria, which may lead to selecting SNPs with spurious association to phenotype. Moreover, because RMLM and EMEB only undergo one iteration, EMEB alone cannot completely remove spuriously associated SNPs. In BLINK, pseudo-QTNs are selected based on BIC values generated by the model, and as a result, associated SNPs close to but not matching the QTN may also appear to have BICs with optimal value.

SEIR effectively reduces the number of SNPs from the original set to facilitate fast variable selection in a reduced data space. The, using a small subset of the reduced set of SNPs rather than the full, original set of SNPs, the embedded selector in SEIR can quickly select pseudo-QTNs with a relatively low computational burden. In addition, new pseudo-QTNs are added to the set of previous covariates as new covariates after each iteration in FarmCPU and BLINK [17, 19]. Thus, a false SNP marker introduced as a covariate in a previous round will be retained in subsequent iterations, resulting in an increased number of false positives and lower statistical power. By contrast, in SEIR, the embedded selector can reserve parsimonious covariates and avoid problems stemming from “only-in-no-out” limitations that prevent removal of wrong or non-significant covariates. This feature of SEIR is likely an important contributing factor to its increased statistical power.

In phenotype simulation studies, we also found that different multi-locus methods can identify an overlapping set of significant SNPs, suggesting their robustness. Interestingly, each multi-locus method also identified some unique significant SNPs, which implied that no single method can capture all trait-associated SNPs. This limitation may be due to the variability of genetic architecture responsible for determining different complex traits. These results align well with previous studies that reported different GWAS methods should be used in combination to improve detection rates and statistical robustness [35, 36].

When applied to real phenotype data from multiple species, SEIR, MLMM, FarmCPU, MRMLM and BLINK could successfully identify previously reported associated SNPs. However, the sets of significant SNPs detected by each method do not completely overlap, reinforcing the findings obtained through phenotype simulation tests. For example, the significant SNPs associated with backfat thickness in pigs identified by each multi-locus method are located within a LD block on chromosome 7, with SEIR identifying an additional, significant SNP in close proximity within the LD block. Multi-omics analyses integrating LD analysis for SNPs with Hi-C, epigenetic and RNA-seq datasets from pig adipose tissue revealed clear biological relationships between the identified SNPs and backfat thickness, thus supporting the reliability of SEIR-based SNP identification and providing a potentially informative and robust approach for characterizing significant SNPs.

## Conclusions

This study proposes the use of the Selector-Embedded Iterative Regression (SEIR) tool to enhance statistical power in multi-locus GWAS. SEIR integrates a multi-functional selector with iterative single-marker scanning to efficiently identify reliable pseudo-QTNs. Single-marker scanning substantially reduces the number of candidate markers, decreasing the computational burden, while the multi-locus model-based selector reduces the FDR and number of Type I errors in selecting pseudo-QTNs as covariates for fixed model analysis, consequently decreasing spuriously associated SNPs and improving statistical power. Proof-of-concept evidence from real and simulated phenotype data demonstrate the statistical robustness, efficiency, and versatility of SEIR as a publicly available GWAS tool for marker detection in studies of complex, multi-locus traits.

## Materials and Methods

### SEIR method

We have developed a SEIR algorithm, which iteratively integrates single-marker scanning with an embedded selector. The single-marker scanning conducted by the Fixed Effect Model (FEM), which tests M markers one at a time, and pseudo-QTNs are included as covariates to control false positives. Specifically, the FEM can be represented as follows:

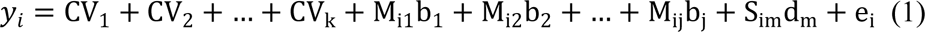

where y_i_ is the observation on the i^th^ individual; CV_1_, CV_2_, …, CV_k_ are other covariates. M_i1_, M_i2_, …, M_ij_ are the genotypes of j pseudo-QTNs, initiated as an empty set; b_1_, b_2_, …, b_j_ are the corresponding effects of the pseudo-QTNs; S_im_ is the genotype of the i^th^ individual and m^th^ genetic marker; d_m_ is the corresponding effect of the m^th^ genetic marker; e_i_ is the residual having a distribution with zero mean and variance of 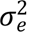.

The goal of FEM is to calculate the P values for all M testing markers. After obtaining the P-value for each marker in the first step, those with P-value larger than the threshold are filtered out (Bonferroni correction, α = 0.01). The remaining SNPs may exhibit linear dependence. If two SNPs have Pearson correlation coefficients above a threshold (0.7), the less significant SNP is removed. This process is repeated until the last SNP is selected, effectively reducing the number of markers to a manageable size. Following the filtering process, the remaining t markers, which are sorted and not highly correlated with each other, are used to select the optimal set of pseudo-QTNs. Using the regression model to represent:

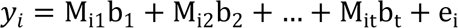

The number of covariate pseudo QTNs is varied to select the optimum set of the j out of t pseudo-QTNs. The optimization is performed using one of the selectors such as stepwise regression [12], smoothly clipped absolute deviation (SCAD) [21], minimax concave penalty (MCP) [22], multivariate adaptive regression splines (MARS) [23], least angle regression (LAR) [24], generalized linear models with L0, L1 penalty [25], among others. Subsequently, the selected k markers are used as the set of pseudo QTNs in equation (1). This process is iterated until the pseudo QTNs remain the same. We have named this alternative solution as the Selector embedded iterative regression (SEIR) method.

### Real data

We used previously published datasets from multiple species, encompassing seven species such as Arabidopsis, human, yeast, mouse, chicken, horse, and pig. Markers with a minor allele frequency of 5% or below were excluded from the original datasets. The principal components were calculated by PLINK using all the SNPs [37]. The Manhattan plots of GWAS results were drawn using CMplot [38].

The human dataset, titled as “East Asian lung cancer dataset”, ID # phs000716.v1. p1, was obtained from dbGaP [28]. In accordance with the privacy and intentions of research participants, the data is only accessible with the permission of NIH (National Institute of Health) and Intramural NCI (National Cancer Institute). The authors acquired the data through dbGaP Authorized Access. A total of 8,807 samples were used, comprising 4,962 lung cancer cases and 3,845 controls. All samples were genotyped, using the Illumina Human610_Quadv1_B and Human660W-Quad_v1_A platforms, with each sample containing 629,968 SNPs.

We used two datasets of Arabidopsis thaliana. The larger dataset, containing 1,179 individuals that were genotyped with 214,545 SNPs, was used for power and FDR simulation tests (Dataset: 2010 project 250K SNP chip genotypes v3.04). The smaller dataset includes 199 samples, with 216,130 SNPs and 107 phenotypes, which were used in GWAS analysis [39].

The mouse dataset was obtained from a heterogeneous stock mice population owned by the Welcome Trust Centre for Human Genetics. This dataset has 1,940 samples with 12,226 SNPs. The real phenotype is the weight growth intercept for GWAS analysis [40].

The chicken dataset has 1,064 samples, with 294,705 SNPs. The real phenotype is egg weight for GWAS analysis [29].

The horse data includes 14 domestic horse breeds and 18 evolutionarily related species. In total, 480 horses were genotyped with a designed ∼ 54,000 SNP assay. A total of 50,621 SNPs was available after removing the SNPs. The trait was coat color for GWAS analysis [41].

The pig genotype dataset includes 4,555 individuals with 47,257 SNPs. The real phenotype is backfat thickness for GWAS analysis.

The yeast dataset contains 4,390 samples, with 28,220 SNPs and 20 growth traits. We use 20 growth traits for GWAS analysis [27].

### Simulated phenotypes

We generated simulated data based on the observed genotypes of the human dataset and Arabidopsis thaliana dataset. Phenotypes were simulated by the simulated phenotype generation function in the Genome Association and Prediction Integrated (GAPIT) Tool [42], which randomly samples *m* SNPs as quantitative trait nucleotides (QTN) and generates the effect of each QTN (*m*) from a standard normal distribution or from a geometric distribution. Subsequently, it calculates the genetic value of each individual as 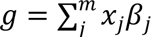, where *x* is coded as 0, 1, and 2 for genotypes *aa*, *Aa*, and and *AA*, respectively. Finally, it generates the residual effects (*e*) from *N*(0, *var*(*g*)(1 − ℎ^2^)/ℎ^2^) and calculates the simulated phenotype as *y* = *g* + *e*.

We used different values for trait heritability (ℎ^2^ = 0.2, 0.6, and 0.8) and for the number of QTN (*m* = 20, 100, 500, and 1000). The effect of each QTN (*m*) was generated from a normal distribution (for *m* = 20) or from a geometric distribution (for *m* = 100, 500 and 1000). For each setting, we repeated the simulation 100 times, randomizing the positions of the QTN in each replicate.

### Power examination under different levels of Type I error and FDR

We used the receiver operating characteristic (ROC) curve to compare diffierent methods for their efficiencies in the detection of significant effiects under different levels of FDR and type I error. A QTN was considered identified if a positive marker was within a prescribed interval distance (e.g., 10 kb). Power was defined as the proportion of QTNs identified at a threshold of Type I error or FDR. Markers were used to derive the null distribution of negative control if no QTN was within the interval. The null distribution of Type I error was derived from the non-QTN markers. FDR was defined as the proportion of the non-QTN markers among the positive markers.

### Hi-C data collection

The Hi-C experiments were performed in fat tissue from two adult LW pigs, flowing in situ ChIA-PET [43] and BL-Hi-C [44] protocols. The fat tissue was fixed with 1% formaldehyde at room temperature for 20 min and then treated with a final concentration of 0.2 M glycine solution to quench the reaction for 10 min at room temperature. The tissue was mixed with 500ul ice-col Hi-C lysis buffer on ice for 20 min. Then the tissue pellet was resuspended in 500ul 0.5% SDS and incubated at 62 °C for 10min. Subsequently, a final 10% Triton X-100 was added at 1% final concentration to quench the SDS and incubated at 37 °C for 10 min. The enzyme digestion and linker ligation followed in situ ChIA-PET [43] and BL-Hi-C [44] protocols as well. The Hi-C DNA was extracted and prepared for sequencing library construction with the Tn5 enzyme from Vazyme Company. Finally, two Hi-C libraries were obtained, one from each individual. The Hi-C libraries were sequenced on an Illumina nova-seq PE150 platform.

### Hi-C data processing

The sequence data were processed using the CUP pipeline [45] with xx parameters. The bam file were further processed with the HiC-Pro [46] pipeline to generate validate pairs. Then, validated pairs from different individuals were merged to build raw contact maps, and ICE normalization was performed on contact maps using the HiC-Pro [46] pipeline as well. Visualized contact matrices were generated by juicer tools [46] (https://github.com/aidenlab/juicer/wiki/Download). The Hi-C loops calling were used HiCCUPS with minor modifications at 25kb resolution (-r 25000) [47].

### Integration analysis pig epigenetics data

To explore the relationship between the LD of SNPs and 3D genome structure, we first obtained whole-genome resequencing data of 163 swine including Duroc, Landrace, Yorkshire from ISWINE database [48]. After data quality control using PLINK [37] (removing SNPs with a missing rate greater than 5%, or a minimum allele frequency of less than 1%, or a p-value of less than 1e-6 in the Hardy-Weinberg equilibrium test), 16,141,439 SNPs were used for analysis. Next, a 1 Mb genomic region was randomly chosen in pig genome. SNPs located in this region were selected randomly per 5kb distance, and the normalized coefficient of linkage disequilibrium (D’) was used to characterize LD within 1Mb distance. The LD heat map of whole-genome resequencing data and Hi-C contact map were built at the 5 kb resolution. D’ was calculated by LDblockshow [49]. The visualization of LD heat map of whole-genome resequencing data was carried out using LDheatmap [50]. The Hi-C heat map was obtained using Juicebox tool. We then calculated the D’ of the whole genome of in our backfat thickness pig population using the Illumina PorcineSNP50 Bead Chip, which includes 47,257 SNPs. In further comparative analysis, we counted the D’ between SNPs within a TAD and the D’ between the SNPs spanning the two vicinity TADs. P-value statistics were performed using rank sum test method. The overlap between LD blocks and TAD regions was determined using intersect command of BEDTools [51].

The ATAC-seq, ChIP-seq (H3K27ac and H3K4me3) and RNA-seq of pig fat tissue were from the previous study [52]. Visualization of Hi-C loops, ATAC-seq, ChIP-seq (H3K27ac and H3K4me3) and RNA-seq of pig fat tissue and LD block harbored SNPs significantly associated backfat thickness was performed using IGV browser [53].

## Declarations

### Ethics approval and consent to participate

No ethical approval was required for this study.

### Consent for publication

Not applicable

### Availability of data and materials

#### Human dataset

The human dataset, titled as “East Asian lung cancer dataset”, ID # phs000716.v1. p1, was obtained from dbGaP (URL: http://www.ncbi.nlm.nih.gov/projects/gap/cgi-bin/study.cgi?study_id=phs000716.v1.p1). In accordance with the privacy and intentions of research participants, the data is only accessible with the permission of NIH (National Institute of Health) and Intramural NCI (National Cancer Institute). The authors acquired the data through dbGaP Authorized Access.

Arabidopsis thaliana dataset: https://github.com/Gregor-Mendel-Institute/atpolydb

#### Mouse dataset

Data are from the “Neves HHR, Carvalheiro R, Queiroz S a. A comparison of statistical methods for genomic selection in a mice population. BMC Genet. 2012;13:100.doi:10.1186/ 1471-2156-13-100” study whose authors may be contacted at Haroldo HR Neves

#### Yeast dataset

https://www.ncbi.nlm.nih.gov/pmc/articles/PMC4635962/bin/ncomms9712-s6.zip Chicken dataset: https://figshare.com/articles/dataset/Genome-wide_Association_Analysis_of_Age-Dependent_Egg_Weights_in_Chickens/5844420 Horse dataset: https://figshare.com/articles/Horse_dataset/12336773

#### Pig dataset

https://doi.org/10.6084/m9.figshare.21130672

## Authors’ contributions

MJZ, GLZ, and YXZ conceived and designed the experiments. GLZ, FJX, and JKQ conducted the simulation and real data analysis. DYW conducted the Hi-C data collection. RZK, LML, and MYH conducted the Hi-C data processing. XLL provided the human data. GLZ, YXZ wrote the first draft of the manuscript, XYL, SHZ provided suggestions for the manuscript. All authors read and approved the final manuscript.

## Acknowledgments

Not applicable

## References

1. Kang HM, Sul JH, Service SK, Zaitlen NA, Kong SY, Freimer NB, et al. Variance component model to account for sample structure in genome-wide association studies. Nat Genet. 2010;42:348–54.

2. Devlin B, Roeder KJB. Genomic control for association studies. Biometrics. 1999;55:997–1004.

3. Pritchard JK, Stephens M, Donnelly P. Inference of population structure using multilocus genotype data. Genetics. 2000;155:945–59.

4. Zhao K, Aranzana MJ, Kim S, Lister C, Shindo C, Tang C, et al. An Arabidopsis example of association mapping in structured samples. PLoS Genet. 2007;3:e4.

5. Yu J, Pressoir G, Briggs WH, Vroh Bi I, Yamasaki M, Doebley JF, et al. A unified mixed-model method for association mapping that accounts for multiple levels of relatedness. Nat Genet. 2006;38:203–8.

6. Pritchard JK, Stephens M, Rosenberg NA, Donnelly P. Association Mapping in Structured Populations. The American Journal of Human Genetics. 2000;67:170–81.

7. Zhang Z, Ersoz E, Lai CQ, Todhunter RJ, Tiwari HK, Gore MA, et al. Mixed linear model approach adapted for genome-wide association studies. Nat Genet. 2010;42:355–60.

8. Svishcheva GR, Axenovich TI, Belonogova NM, van Duijn CM, Aulchenko YS. Rapid variance components-based method for whole-genome association analysis. Nat Genet. 2012;44:1166–70.

9. Loh PR, Tucker G, Bulik-Sullivan BK, Vilhjalmsson BJ, Finucane HK, Salem RM, et al. Efficient Bayesian mixed-model analysis increases association power in large cohorts. Nat Genet. 2015;47:284–90.

10. Jiang L, Zheng Z, Qi T, Kemper KE, Wray NR, Visscher PM, et al. A resource-efficient tool for mixed model association analysis of large-scale data. Nat Genet. 2019;51:1749–55.

11. Yi N, Xu S. Bayesian LASSO for quantitative trait loci mapping. Genetics. 2008;179:1045–55.

12. Cordell HJ, Clayton DG. A unified stepwise regression procedure for evaluating the relative effects of polymorphisms within a gene using case/control or family data: application to HLA in type 1 diabetes. Am J Hum Genet. 2002;70:124–41.

13. Ayers KL, Cordell HJ. SNP selection in genome-wide and candidate gene studies via penalized logistic regression. Genet Epidemiol. 2010;34:879–91.

14. Lu HY, Liu XF, Wei SP, Zhang YM. Epistatic association mapping in homozygous crop cultivars. PLoS One. 2011;6:e17773.

15. Chiaromonte F, Martinelli J. Dimension reduction strategies for analyzing global gene expression data with a response. Mathematical Biosciences. 2002;176:123–44.

16. Segura V, Vilhjalmsson BJ, Platt A, Korte A, Seren U, Long Q, et al. An efficient multi-locus mixed-model approach for genome-wide association studies in structured populations. Nat Genet. 2012;44:825–30.

17. Liu X, Huang M, Fan B, Buckler ES, Zhang Z. Iterative Usage of Fixed and Random Effect Models for Powerful and Efficient Genome-Wide Association Studies. PLoS Genet. 2016;12:e1005767.

18. Wang SB, Feng JY, Ren WL, Huang B, Zhou L, Wen YJ, et al. Improving power and accuracy of genome-wide association studies via a multi-locus mixed linear model methodology. Sci Rep. 2016;6:19444.

19. Huang M, Liu X, Zhou Y, Summers RM, Zhang Z. BLINK: a package for the next level of genome-wide association studies with both individuals and markers in the millions. Gigascience. 2019;8.

20. Wen C, Zhang A, Quan S, Wang X. BeSS: An R Package for Best Subset Selection in Linear, Logistic and Cox Proportional Hazards Models. Journal of Statistical Software. 2020;94:1–24.

21. Fan J, Li R. Variable selection via nonconcave penalized likelihood and its oracle properties. Journal of the American Statistical Association. 2001;67:301–20.

22. Zhang C-H. Nearly unbiased variable selection under minimax concave penalty. The Annals of Statistics. 2010;38.

23. Friedman JH. Multivariate Adaptive Regression Splines. Annals of Statistics. 1991;19:1–67.

24. Efron B, Hastie T, Johnstone I, Tibshirani R. Least Angle Regression. Annals of Statistics. 2003.

25. Tibshirani R. Regression Shrinkage and Selection Via the Lasso. Journal of the Royal Statistical Society: Series B (Methodological). 1996;58:267–88.

26. Chen Y, Yang Y. The One Standard Error Rule for Model Selection: Does It Work? Stats. 2021;4:868–92.

27. Bloom JS, Kotenko I, Sadhu MJ, Treusch S, Albert FW, Kruglyak L. Genetic interactions contribute less than additive effects to quantitative trait variation in yeast. Nat Commun. 2015;6:8712.

28. Lan Q, Hsiung CA, Matsuo K, Hong YC, Seow A, Wang Z, et al. Genome-wide association analysis identifies new lung cancer susceptibility loci in never-smoking women in Asia. Nat Genet. 2012;44:1330–5.

29. Liu Z, Sun C, Yan Y, Li G, Wu G, Liu A, et al. Genome-Wide Association Analysis of Age-Dependent Egg Weights in Chickens. Front Genet. 2018;9:128.

30. McCue ME, Bannasch DL, Petersen JL, Gurr J, Bailey E, Binns MM, et al. A High Density SNP Array for the Domestic Horse and Extant Perissodactyla: Utility for Association Mapping, Genetic Diversity, and Phylogeny Studies. PLOS Genetics. 2012;8:e1002451.

31. Michaels SD, Amasino RM. FLOWERING LOCUS C encodes a novel MADS domain protein that acts as a repressor of flowering. The Plant Cell. 1999;11:949–56.

32. Pahl R, Schäfer H. PERMORY: an LD-exploiting permutation test algorithm for powerful genome-wide association testing. Bioinformatics. 2010;26:2093–100.

33. Claussnitzer M, Dankel SN, Kim KH, Quon G, Meuleman W, Haugen C, et al. FTO Obesity Variant Circuitry and Adipocyte Browning in Humans. N Engl J Med. 2015;373:895–907.

34. Ricoult SJ, Manning BD. The multifaceted role of mTORC1 in the control of lipid metabolism. EMBO Rep. 2013;14:242–51.

35. Zhang Y-M, Jia Z, Dunwell JM. Editorial: The Applications of New Multi-Locus GWAS Methodologies in the Genetic Dissection of Complex Traits. 2019;10.

36. Zhou G-L, Xu F-J, Qiao J-K, Che Z-X, Xiang T, Liu X-L, et al. E-GWAS: an ensemble-like GWAS strategy that provides effective control over false positive rates without decreasing true positives. Genetics Selection Evolution. 2023;55:46.

37. Purcell S, Neale B, Todd-Brown K, Thomas L, Ferreira MA, Bender D, et al. PLINK: a tool set for whole-genome association and population-based linkage analyses. Am J Hum Genet. 2007;81:559–75.

38. Yin L, Zhang H, Tang Z, Xu J, Yin D, Zhang Z, et al. rMVP: A memory-efficient, visualization-enhanced, and parallel-accelerated tool for genome-wide association study. Genomics Proteomics Bioinformatics. 2021;19:619–28.

39. Atwell S, Huang YS, Vilhjálmsson BJ, Willems G, Horton M, Li Y, et al. Genome-wide association study of 107 phenotypes in Arabidopsis thaliana inbred lines. Nature. 2010;465:627–31.

40. Neves HH, Carvaheiro R, Queiroz SA. A comparison of statistical methods for genomic selection in a mice population. BMC Genet. 2012;2012:13:100.

41. McCue ME, Bannasch DL, Petersen JL, Gurr J, Bailey E, Binns MM, et al. A high density SNP array for the domestic horse and extant Perissodactyla: utility for association mapping, genetic diversity, and phylogeny studies. PLoS Genet. 2012;8:e1002451.

42. Wang J, Zhang Z. GAPIT Version 3: boosting power and accuracy for genomic association and prediction. Genomics Proteomics Bioinformatics. 2021;19:629–40.

43. Wang P, Feng Y, Zhu K, Chai H, Chang YT, Yang X, et al. In situ Chromatin Interaction Analysis Using Paired-End Tag Sequencing. Curr Protoc. 2021;1:e174.

44. Liang Z, Li G, Wang Z, Djekidel MN, Li Y, Qian MP, et al. BL-Hi-C is an efficient and sensitive approach for capturing structural and regulatory chromatin interactions. Nat Commun. 2017;8:1622.

45. Bertolini JA, Favaro R, Zhu Y, Pagin M, Ngan CY, Wong CH, et al. Mapping the Global Chromatin Connectivity Network for Sox2 Function in Neural Stem Cell Maintenance. Cell Stem Cell. 2019;24:462–76 e6.

46. Servant N, Varoquaux N, Lajoie BR, Viara E, Chen CJ, Vert JP, et al. HiC-Pro: an optimized and flexible pipeline for Hi-C data processing. Genome Biol. 2015;16:259.

47. Rao SS, Huntley MH, Durand NC, Stamenova EK, Bochkov ID, Robinson JT, et al. A 3D map of the human genome at kilobase resolution reveals principles of chromatin looping. Cell. 2014;159:1665–80.

48. Fu Y, Xu J, Tang Z, Wang L, Yin D, Fan Y, et al. A gene prioritization method based on a swine multi-omics knowledgebase and a deep learning model. Commun Biol. 2020;3:502.

49. Dong SS, He WM, Ji JJ, Zhang C, Guo Y, Yang TL. LDBlockShow: a fast and convenient tool for visualizing linkage disequilibrium and haplotype blocks based on variant call format files. Brief Bioinform. 2021;22.

50. Ji-Hyung Shin, Sigal Blay, Brad McNeney, Graham J. LDheatmap: An R Function for Graphical Display of Pairwise Linkage Disequilibria between Single Nucleotide Polymorphisms. Journal of Statistical Software. 2006;16.

51. Quinlan AR, Hall IM. BEDTools: a flexible suite of utilities for comparing genomic features. Bioinformatics. 2010;26:841–2.

52. Zhao Y, Hou Y, Xu Y, Luan Y, Zhou H, Qi X, et al. A compendium and comparative epigenomics analysis of cis-regulatory elements in the pig genome. Nat Commun. 2021;12:2217.

53. Robinson JT, Thorvaldsdóttir H, Winckler W, Guttman M, Lander ES, Getz G, et al. Integrative genomics viewer. Nature Biotechnology. 2011;29:24–6.

